# Deletion of the gene for the African swine fever virus BCL-2 family member A179L increases virus uptake and apoptosis, but decreases virus spread in macrophages and reduces virulence in pigs

**DOI:** 10.1101/2023.04.27.538639

**Authors:** Ana Luisa Reis, Anusyah Rathakrishnan, Leah V. Goulding, Claire Barber, Lynnette C. Goatley, Linda K. Dixon

## Abstract

African swine fever virus encodes proteins that inhibit apoptosis including one member of the BCL-2 family, A179L. Deletion of the A179L gene from the virulent genotype I isolate Benin 97/1 compared to Benin 97/1 expressing A179L or mock-infected macrophages, resulted in increased Caspase 3 and 7 activity, annexin V binding to surface phosphatidyl serine and DNA fragmentation, measured by terminal deoxynucleotidyl transferase nick-end labelling. These results confirmed that apoptosis was induced earlier in macrophages infected with the BeninΔA179L virus. Increased cell entry of the A179L gene-deleted virus was indicated at early times since up to double the numbers of cells expressed fluorescent protein from the virus genome. Yields of infectious virus were similar over a single cycle but were significantly lower for the A179L gene-deleted virus over a multi-step growth cycle. Pigs immunised and boosted with the BeninΔA179L virus showed no clinical signs, although a weak cellular response to ASFV was observed showing that the virus had replicated. The immunised pigs were not protected against challenge with the virulent parental virus Benin 97/1 although viremia was lower at 3 days post-challenge compared to the control non-immune pigs. The reduced levels of virus replication in macrophages probably limited induction of a protective immune response. The results show an important role for the A179L protein in virus replication in macrophages and virulence in pigs.

**IMPORTANCE:** African swine fever virus (ASFV) causes a lethal disease of pigs that has spread extensively in Africa, Europe and Asia. The virus codes for more than 150 proteins, many of which help the virus to evade the host’s defences following infection. We investigated the effect of deleting one of these genes, A179L, from the genome of an ASFV isolate that causes death of infected pigs. A179L belongs to the BCL-2 protein family, consisting of members which promote or inhibit apoptosis with A179L belonging to the latter. Deleting the A179L gene reduced ASFV replication and spread between macrophages, its main target cells. This was correlated with an increase in cell death. Pigs infected with the virus with A179L gene deleted did not show signs of disease and no virus replication was detected in blood. A low immune response was generated but the immunised pigs were not protected when challenged with the parental deadly virus. The results show that the A179L gene is important for ASFV to replicate efficiently in cells and in animals.

---

African swine fever (ASF) is a highly contagious disease of domestic pigs and wild boar caused by a double-stranded DNA virus, African swine fever virus (ASFV) (1, 2). ASFV is the only member of the *Asfarviridae* family and replicates in the cytoplasm of cells of the myeloid lineage, primarily macrophages or monocytes at an intermediate or late stage of differentiation (3). In East Africa, ASFV is maintained in a sylvatic cycle involving warthogs (*Phacochoerus africanus)* and soft ticks of the genus *Ornithodoros* (4). The virus can persist in warthogs and bushpigs with few clinical signs of disease, but case fatality rates in domestic pigs or wild boar infected with virulent isolates of ASFV can approach 100% (5-8). In 2007 ASFV was introduced to the Republic of Georgia (9) and spread to Russia, Eastern Europe, and EU member countries. Following the introduction to China in 2018 disease spread extensively there and in other countries in Southeast Asia causing high economic losses (10-12). Vaccines are not widely available, thereby limiting control of ASFV outbreaks. Ongoing efforts centre on the deletion of target genes to construct modified live-attenuated vaccines (13-17).

The ASFV genome varies between 170 to 193 kbp and encodes up to 170 proteins. Many of the virus genes are not essential for replication but have important roles in modulating the host’s antiviral defences. Amongst these are inhibitors of the type I interferon response and inhibitors of apoptosis (1, 18). Apoptosis of infected host cells is an effective protective mechanism to disrupt virus replication and limit the spread of progeny virus (19). ASFV encodes several regulators of host apoptotic pathways including; an IAP homolog (A224L) that inhibits caspase 3 activity, an inhibitor of the pro-apoptotic stress-induced CHOP protein dependent pathway (DP71L), a viral C-type lectin that inhibits p53 activity (EP153R) and a viral BCL-2 protein family member (A179L) (20-23).

The BCL-2 family proteins regulate the host cell’s commitment to apoptosis through interplay between the BCL-2 family members. Ultimately mitochondrial outer membrane (MOM) permeabilization is regulated via pores formed by pro-apoptotic BCL-2 family proteins and subsequent release of apoptosis mediators into the cytosol. The BCL-2 family members are broadly divided into the anti-apoptotic or ‘pro-survival’ members including BCL-2, BCL-x_L_, MCL-1, BCL-W and BFL-1/A1(24, 25). The pro-apoptotic members include pore forming proteins (BAX, BAK and BOK) and the BH3-only proteins (such as BAD, BID and BIK) (26, 27). The BH3-only proteins function either by binding and ‘activating’ the pore forming proteins or through binding to anti-apoptotic proteins and ‘freeing’ sequestered pro-apoptotic regulators (27, 28). Anti-apoptotic BCL-2 proteins sequester the pro-apoptotic family members to prevent permeabilization of the mitochondrial membrane (24, 25). Several viruses code for homologs of the anti-apoptotic BCL-2 family members to regulate apoptosis (29-31).

The ASFV A179L protein has anti-apoptotic activity when expressed exogenously in cells and is proposed to promote cell survival to facilitate ASFV replication (21, 32), although until now this activity has not been demonstrated in ASFV infected cells. A179L contains 3 BCL-2 homology domains (BH4, BH1 and BH2) and localizes to the mitochondria or ER (33, 34). The protein associates with pro-apoptotic BCL-2 proteins (such as Bid, Bim and Bad) with varying affinities, in addition to autophagy associated protein Beclin-1 (35-37). A179L, like the anti-apoptotic BCL-2 family members such as BCL-2 and BCL-x_L_, may inhibit apoptosis through association with the BH3 motifs of the pro-apoptotic regulators, thereby interfering with their activity and inhibiting cell death to extend virus replication (36).

To further understand the role of the A179L protein during ASFV infection, we inactivated the A179L gene on the genome of the virulent ASFV isolate Benin 97/1. This resulted in a reduction in Benin 97/1 replication in porcine primary macrophages by 10-fold over a multistep growth curve. Increases in activation of caspases 3 and 7, DNA fragmentation measured by Terminal deoxynucleotidyl transferase nick-end labelling (TUNEL) staining and levels of Annexin V surface binding were observed from early times post-infection following infection with the BeninΔA179L deletion mutant when compared to the parental virulent virus. These results support the conclusion that A179L suppresses caspase activation and apoptosis in infected macrophages. Live cell imaging of macrophages infected with wild type or A179L gene deleted viruses, expressing fluorescent proteins, showed a much greater infection rate by the A179L deletion mutant. This may result from incorporation of phosphatidyl serine into the external envelope and uptake mediated by phosphatidyl serine receptors. We also showed that the deletion of the A179L gene attenuated the Benin 97/1 virus, since no clinical signs or viremia were observed following infection. However, the pigs were not protected against challenge with the parental virulent virus suggesting insufficient virus replication had occurred to induce a protective response. The results show that A179L has an important role in ASFV replication in macrophages and virulence in pigs.

## RESULTS

### Generation of A179L recombinant viruses

To delete the A179L gene from Benin 97/1, a transfer vector pcDNA3_A179L-LF_VP72-GUS_A179L-RF was constructed to include the left and right flanking regions of A179L from the ASFV genome. The β-glucuronidase (GUS) reporter gene, under control of the VP72 (B646L) gene promoter was inserted between the flanking regions replacing sequences from the A179L gene (Figure 1A). A179L is read towards the left genome end and is separated by a single base between the stop codon of A179L and the start codon of A859L. We therefore incorporated 157 bp of the 3’ end of A179L gene into the left flank (LF) of the transfer vector, so that the promoter of A859L would not be interrupted. The A859L gene is expressed late during infection and its promoter region may be within the 3’ end of the A179L coding region which is expressed early (38). A859L codes for a protein belonging to helicase superfamily 2. Deletion of this gene from a virulent ASFV isolate, Georgia/2007, did not reduce virus replication in macrophages or virulence in pigs (39) so any changes in its expression would not be expected to affect virus growth. The 157 bp region of A179L included in the transfer vector contains the sequence that encodes residues at the C-terminus of A179L from 129 to 179 amino acids. However, this is unlikely to be expressed as there is no in-frame upstream translation start codon. The 157 bp fragment contains sequences coding for most of the C-terminal BCL-2 homology domain 2 (BH2) and lacks just the two N-terminal residues from this domain. However, the upstream BH4 and BH1 domains are not present in the 157 bp fragment remaining from the A179L gene. Structural analysis of A179L bound to BH3 domains from BID and BAX showed that residues encompassing all 3 BH domains of A179L between residues 2 to 141 are involved in binding these pro-apoptotic proteins. Specific residues including R86, G76 and Y86 are important for the binding to BH3 domain proteins. Therefore, in the unlikely event that this fragment remaining at the C-terminus of A179L was expressed it is unlikely to be functional. The transfer vector was transfected into porcine alveolar macrophages (PAMs) infected with Benin 97/1 virus and recombinant virus BeninΔA179L was identified by expression of the reporter gene in infected cells and purified by limiting dilution over 10 passages. Two additional recombinant viruses were constructed each expressing an mNeonGreen fluorescent protein in order to follow infections of macrophages by live cell imaging. In one of these, the A179L gene was deleted from the Benin97/1 virus using the same flanking regions as for the GUS expressing mutant virus, but using mNeonGreen under control of the ASFV VP30 (CP204L) gene promoter as reporter to isolate recombinant viruses by single cell sorting (40). This recombinant virus is referred to as BeninΔA179L_mNG (Figure 1A). To provide a control virus, the mNeonGreen reporter was inserted at position 15358 in a non-coding area of Benin 97/1 genome between MGF300-1L and MGF300-2R avoiding disruption of other ORFs in the region (pNG3, pNG1, J64R) (41). This recombinant virus is referred to as Benin_mNG. These virus genome structures were confirmed by PCR and sequence analysis across the sites of the insertion and deletion.

**Figure 1.**
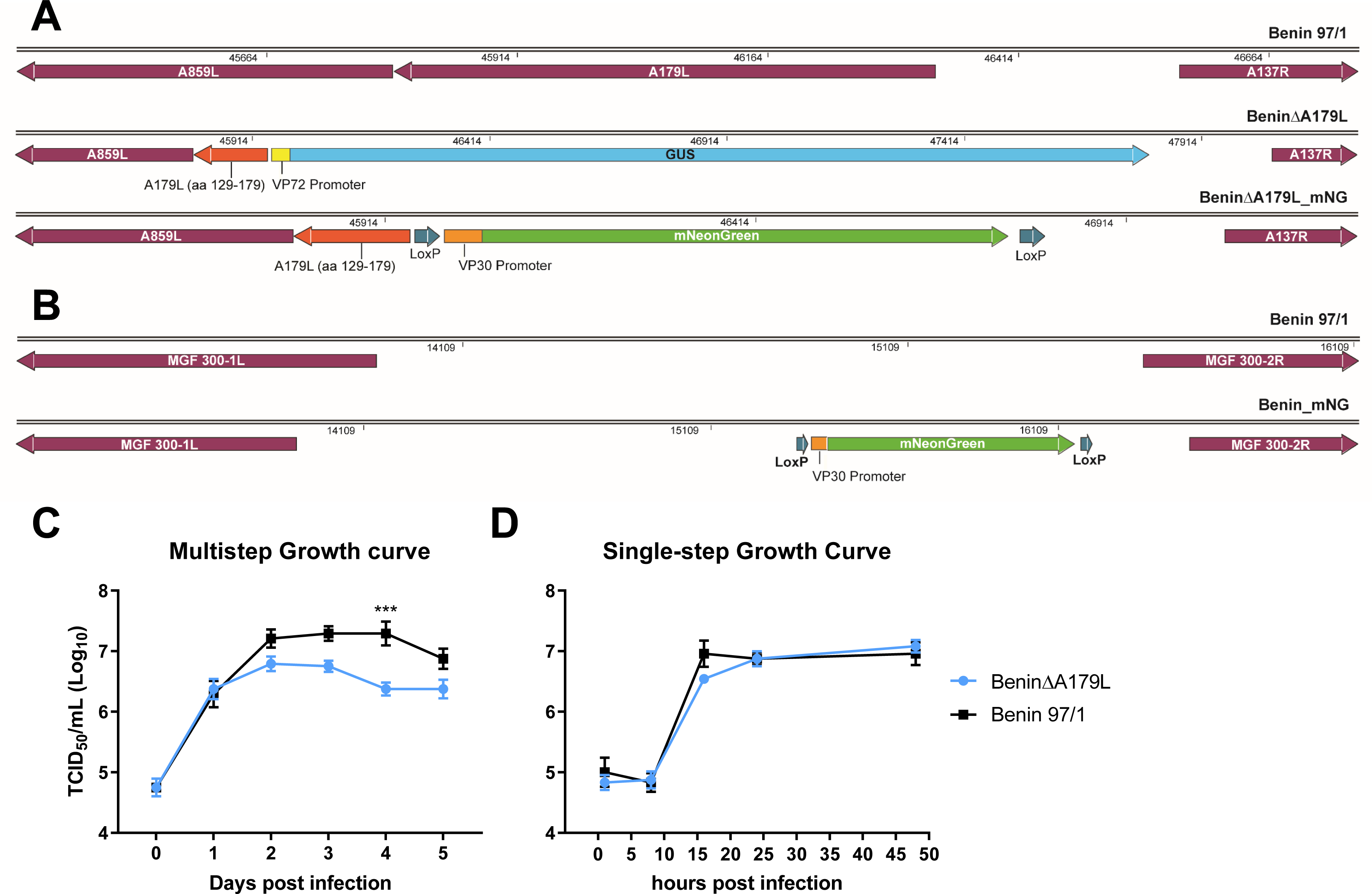
Deletion of A179L gene from ASFV Benin97/1 strain. (**A**) A schematic diagram depicting the deletion of A179L from Benin 97/1 ASFV isolate is shown. In the top panel the position of the A179L gene and flanking genes A859L and A137R are shown. In the middle panel, a diagram of the genome with A179L gene deleted and the β-GUS reporter gene under control of the ASFV p72 promoter inserted in the genome is shown. Note that a fragment of the A179L gene (coding for residues 129 to 179) is retained in the genome to avoid disrupting the promoter for the downstream A859L gene. In the bottom panel the same A179L gene deletion is shown and instead of β-GUS reporter gene, a fluorescent reporter gene, mNeonGreen, under control of the ASFV p30 promoter was inserted. (**B**) The mNeonGreen gene under control of ASFV p30 promoter was inserted in the non-coding region of Benin 97/1 at position 15358 between genes MGF300-1L and MGF300-2R to produce a fluorescent-tagged Benin 97/1. Purified PBMs from 2 different pigs were infected with viruses at MOI 0.01 in triplicates over 5 days (**C**) or at MOI 1.0 for 48h (**D**). Viruses were harvested from both cells and supernatants at different time points indicated (x-axis) and titrated on Vero or PBMs cells in quadruplicates (y-axis). Day 0 shows virus in the inoculum (**C-D**). 2-way ANOVA with Šídák’s multiple comparisons test was performed to evaluate the differences between the recombinant viruses and the wildtype virus. Significant differences are represented by asterisk where *** is p<0.001.

### Complete genome sequencing of recombinant virus BeninΔA179L

The complete genome sequence of the first recombinant virus constructed, BeninΔA179L, confirmed the deletion of the A179L gene and insertion of the β−Glucuronidase reporter construct in its place as expected. We also detected 2 SNPs: the first, a single nucleotide deletion in the non-coding region after A179L (position 50231 on AM712239) and the second, a single nucleotide deletion in the EP402R/CD2v gene at position 69854 (AM712239). This was present at high frequency in the sequence reads (98.6%), induced a predicted frame shift in the coding sequence of the EP402R/CD2v protein at residue 267 and introduced an in-frame termination codon in a different reading frame, 155 residues downstream. In contrast, the A179L gene-deleted virus which expresses the mNeonGreen reporter (BeninΔA179L_mN) was confirmed, by Sanger sequencing of PCR fragments generated from the EP402R/CD2v gene region, to have an identical EP402R/CD2v gene sequence to the parental Benin97/1 isolate.

### Replication of BeninΔA179L deletion mutant viruses in cells

The replication kinetics of the BeninΔA179L virus in porcine bone marrow cells (PBMs) were compared with that of parental Benin 97/1 virus over 5 days using a low multiplicity of infection (MOI 0.01). Virus was harvested from the combined supernatants and cell pellets and titrated. The results showed (Figure 1C) that similar virus titres (10^6.3^ TCID_50/ml_) were obtained at day 1 post-infection. However, from day 2 post-infection consistently lower titres were obtained from cells infected with BeninΔA179L. This difference was statistically significant at day 4 (p=0.0004) with titres of 10^6.4^ TCID_50_/ml and 10^7.3^ TCID_50_/ml for BeninΔA179L and Benin 97/1 respectively. Thus, deletion of the A179L from Benin 97/1 impaired virus replication in PBMs over a multi-step growth curve. A similar reduction in yield of virus BeninΔA179L_mNG was observed over a multistep growth curve (data not shown). This result confirmed that the frame shift mutation we observed in the cytoplasmic tail domain of the BeninΔA179L EP402R/CD2v gene was not the cause of the growth reduction. This was as expected, since deletion of the EP402R/CD2v gene from virulent genotype II and genotype VIII genomes did not reduce virus replication in macrophages or virulence in pigs (42, 43). To determine if replication was also impaired over a single replication cycle, PBMs were infected at a high MOI (1.0) with BeninΔA179L or wild type Benin97/1 and virus was titrated from combined supernatants and cells at different times (1-, 8-, 16-, 24- and 48-hours post-infection). The results (Figure 1D) showed no statistical difference between virus yields obtained at these time points. As expected, virus titres reached a plateau by 16 hours post-infection and these levels were maintained over the course of the experiment (approx. 10^6.8^ TCID_50_/ml). Thus, over a single cycle of replication the total yield of the BeninΔA179L virus grown in PBMs was very similar to that of the parental virulent Benin 97/1 virus.

### Caspase 3/7 activation, in macrophages infected with wild type Benin97/1 and BeninΔA179L viruses

The anti-apoptotic activity of the exogenously expressed ASFV A179L protein has been established, but the impact of this protein during infection of cells was not known (21). Therefore, we examined the difference between the wild type virus and the A179L deletion mutant on apoptotic signalling in infected PBMs, determined by caspase 3/7 activity. A179L is expressed at early times post-infection (38) and therefore we measured caspase 3/7 activity from 2 to 8 hours post-infection. PBMs were mock-infected or infected with Benin 97/1 or BeninΔA179L viruses at a MOI of 0.25. The cells were loaded with 2 µM CellEvent™ Caspase-3/7 Green Detection Reagent for 1 hour prior to infection and fluorescence was measured in live cells every 2 hours until 8 hours post-infection (Figure 2A). There was no significant difference in caspase 3/7 activity in wild type Benin 97/1 infected relative to the mock-infected PBM cells. In contrast, caspase 3/7 activity was significantly increased in BeninΔA179L cells from 4 hours post-infection relative to mock-infected cells (Figure 2A). Early expression of A179L, in cells infected with wild type virus, probably contributed to the reduction of caspase activity to levels similar to mock-infected cells. This contrasted with infections with BeninΔA179L in which increased caspase activation was observed between 4 and 8 hpi. In two of three experiments we observed a small dip in caspase activity at 6 hpi compared to other time points in both mock-infected as well as infected cells (not shown). This dip was not observed when we repeated the experiment using PBM cells that were purified on histopaque gradients suggesting that factors present in the unpurified PBMs may have reduced caspase 3 activity in both infected and uninfected cells.

**Figure 2.**
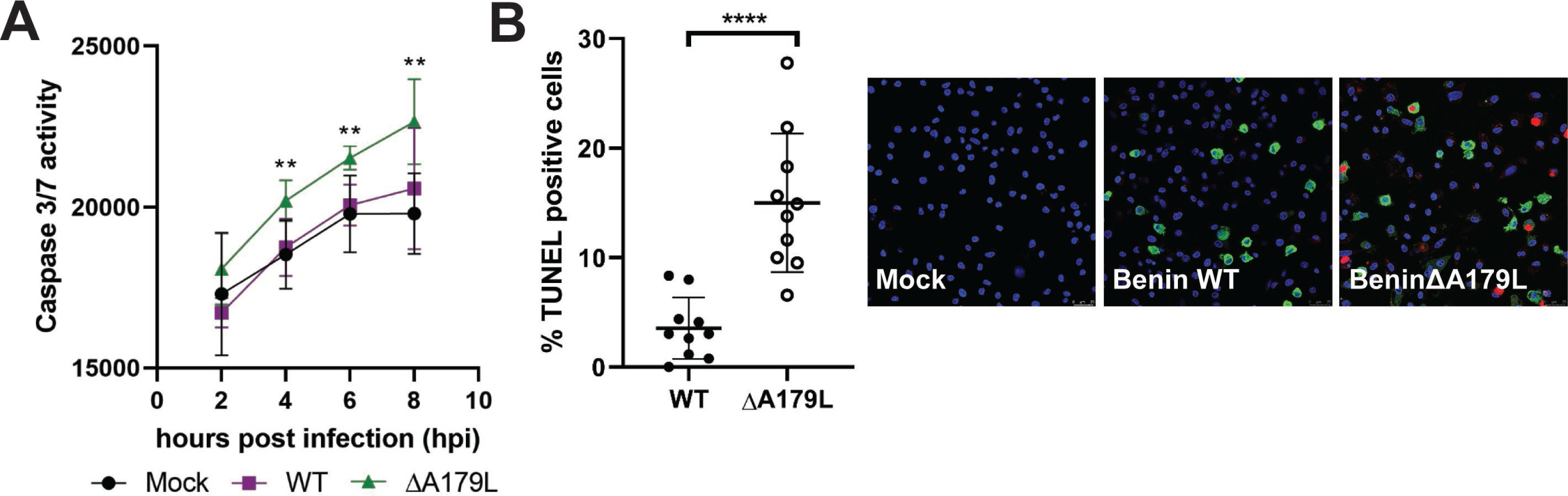
Increased pro-apoptotic signalling in cells infected with BeninΔA179L. (**A**) PBMs were infected with Benin 97/1 or BeninΔA179L at 0.5 MOI. The cells were loaded with 2 μM CellEvent™ Caspase-3/7 Green Detection Reagent after which fluorescence measurements, indicative of caspase3/7 activity, were taken 2, 4, 6, 8 hours post infection (h.p.i.). Significance was determined by 2-way ANOVA between groups within each time point relative to the mock-infected cells. (**B**) PBMs were infected with Benin or BeninΔA179L at 0.25 MOI for 16 h. The cells were stained for TUNEL (red) and ASFV VP30 (green). Cell nuclei labelled with Hoechst 33342 (blue). Representative images displayed. Scale bar 25 µm. Percentage TUNEL positive cells calculated/field of view. Ten fields of view were counted for each virus. The significance was determined by paired t test. Data are expressed as mean values ± SD.

### Deletion of A179L gene results in increased DNA fragmentation measured by TUNEL assay

To provide further evidence that greater levels of apoptosis are induced in cells infected with BeninΔA179L, we used an additional assay, DNA fragmentation by terminal deoxynucleotidyl transferase dUTP nick end labelling (TUNEL) staining. This assay detects apoptotic cells at a late stage. At late times, 16 hours post-infection, the number of apoptotic cells was significantly greater in cells infected with BeninΔA179L compared to cells infected with wild type Benin or mock-infected cells (Figure 2B).

### Deletion of A179L gene results in increased cell surface binding of Annexin V

Exposure of plasma membrane phosphatidylserine during apoptosis allows the binding of Annexin V dye. To confirm that deletion of the A179L gene also results in increased activation of apoptosis, as measured by binding of Annexin V to cell surfaces, we used live cell imaging over an extended time course with the IncuCyte S3. Purified PBM cells were infected at a low MOI (0.1) with wild type or A179L gene-deleted virus each expressing the mNeonGreen reporter under control of the ASFV early P30 promoter (BeninΔA179L_mNG or Benin 97/1_mNG), Binding of Annexin V red fluorescent reagent to live cell surfaces was visualised every 2 h for 48 h. In parallel, infected cells were monitored by expression of mNeonGreen.

Increased Annexin V binding to BeninΔA179L_mNG infected cells was detected from early times, about 4 to 6 hpi, reaching a peak by 16 to 20 hpi and maintaining a plateau or gradually declining after that time point. In contrast, in the control Benin 97/1_mNG infected cells a lower amount of Annexin V binding was observed increasing much more gradually after infection and continuing to rise during the 48 h culture period (Figure 3A). As a positive control for induction of apoptosis, ABT-263 treated cells were monitored in parallel. ABT-263 acts as a mimetic of BH3 domains and inhibits anti-apoptotic members of the BCL-2 family including BCL-2, BCL-xL and BCL-W. In these treated wells, as expected, we also observed a substantial increase in Annexin V levels from 4 hpi (Figure 3A).

**Figure 3.**
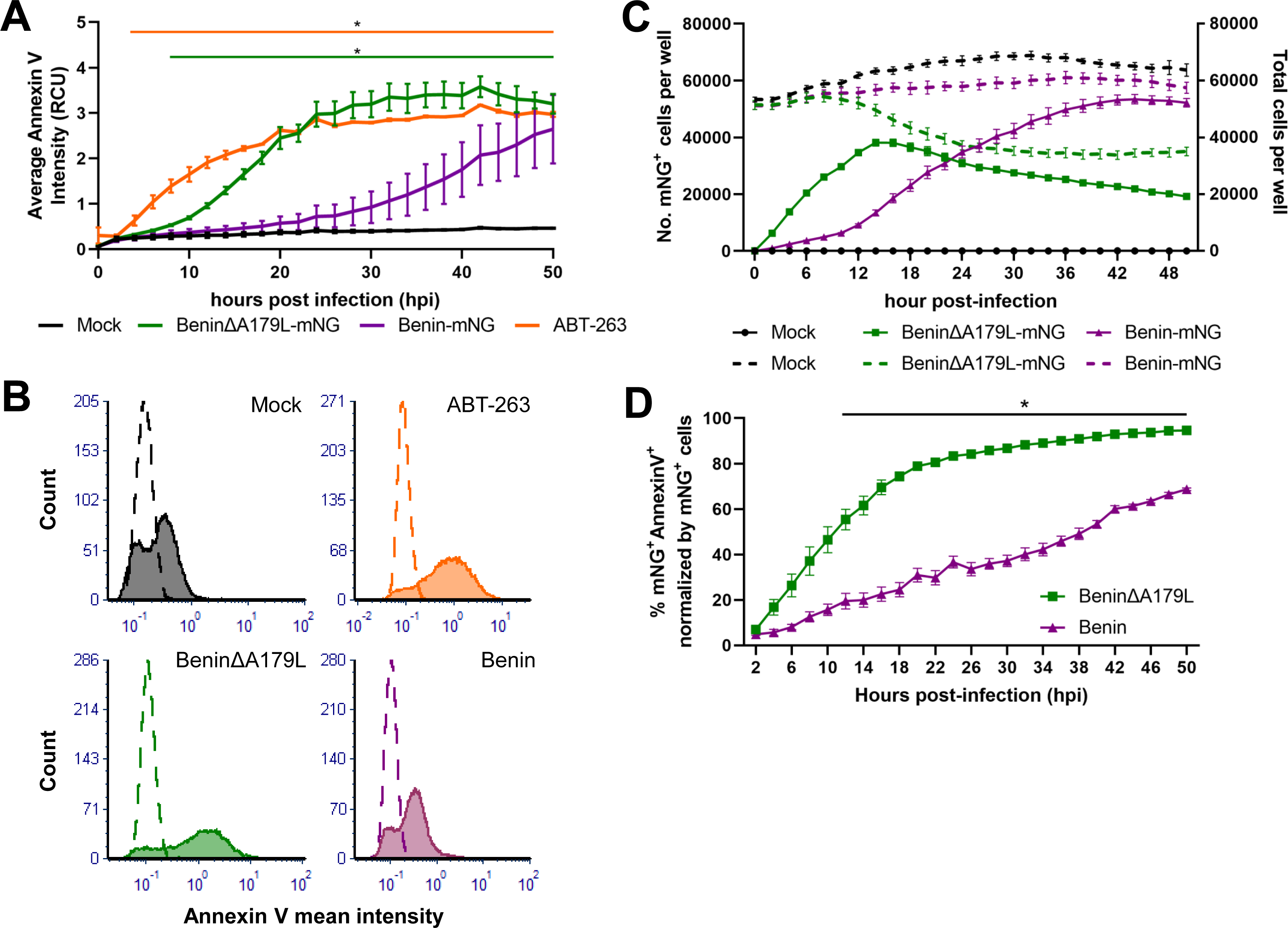
Cell surface Annexin V is induced at early times post-infection with gene-deleted BeninΔA179L_mNG compared to Benin_mNG. Purified PBMs were infected with Benin_mNG or BeninΔA179L_mNG at 0.1 MOI. Annexin V was added onto cells immediately and cells were monitored on the IncuCyte S3 live cell imaging system, imaged every 2h for 48h. Using the adherent cell module, the average Annexin V intensity over time (A), was measured purple lines show Benin-mNG, green shows BeninΔA179L_mNG and orange control cells treated with ABT-263 apoptosis inducer. Panel B shows the mean Annexin V intensity at 22 hpi for mock-infected, ABT-263 treated, Benin_mNG or BeninΔA179L_mNG infected cells. Panel C shows the number of cells expressing mNeonGreen (solid lines) and total number of cells (dashed lines) in green for BeninΔA179L_mNG and purple for Benin_mNG infected cultures. Panel (D) shows the percentage of infected, apoptotic cells (mNeonGreen^+^Annexin^+^). 2-way ANOVA was used to evaluate the differences between groups over time.

Figure 3A shows average Annexin V intensity per well. This may represent increased number of Annexin V positive cells in each well and/or increased Annexin fluorescent intensity in the positive cells. To better understand this finding, the Annexin V mean fluorescent intensity (MFI) per cell was examined at 22hpi, when the ratio of Annexin V levels between BeninΔA179L_mNG and Benin97/1_mNG infected wells was the greatest. Annexin V binding intensity in PBMs infected with Benin97/1_mNG was comparable to that of the mock-infected cells, whereas a dramatic shift in MFI was observed in cells infected with BeninΔA179L (Figure 3B). In fact, this shift was similar to the one observed in the apoptosis-inducer ABT-263 treated PBMs (Figure 3B).

The number of BeninΔA179L_mNG infected cells, assessed by expression of mNG, increased rapidly after infection and reached a peak by 16 hpi thereafter declining gradually. In contrast, numbers of Benin97/1_mNG infected cells increased gradually. By 16 hpi numbers of BeninΔA179L_mNG infected cells were about double those of Benin 97/1_mNG. By 22 to 24 hpi numbers were similar in both infections but numbers of Benin97/1_mNG infected cells continued to rise and reached a plateau by about 36 hpi (Figure 3C, solid lines). The results indicate an increase in BeninΔA179L_mNG virus uptake and early gene expression in cells. One possibility is this may be mediated by binding of phosphatidyl serine, incorporated into the virus external envelope, to receptors on the host cell. The virus growth curves in Figure 1C indicate similar levels of virus replication up to 1-day post-infection, thus the increased numbers of mNeonGreen expressing cells observed in BeninΔA179L_mNG infected cultures post-infection do not apparently result in an increase in infectious virus progeny, perhaps due to abortive infection caused by earlier induction of apoptosis. We also compared total numbers of cells in cultures to determine if infection with BeninΔA179L_mNG caused a depletion in cell numbers (Figure 3C, broken lines). This showed similar numbers of cells in cultures mock-infected or infected with either virus until about 12 hpi and after that, numbers of cells in mock-infected or Benin97/1_mNG remained relatively constant whereas the numbers in the BeninΔA179L_mNG cultures decreased by about 25% at 24 hpi and then remained constant. The decrease in numbers of cells infected with BeninΔA179L_mNG was closely correlated with the number of cells remaining in culture indicating that cell death limited virus replication.

We also evaluated the percentage of mNeonGreen expressing cells that bound to Annexin V to estimate the proportion of cells that were both infected, and apoptotic. As shown in Figure 3D, the percentage of the infected and Annexin V bound cells was also higher following infection with BeninΔA179L_mNG compared to Benin97/1_mNG. This was observed from 4 hpi, although the greatest difference in the percentage of apoptotic cells in cultures infected with these viruses was from about 12 hpi (Figure 3D).

### Immunisation of pigs with BeninΔA179L and challenge with virulent Benin97/1. Scoring of clinical signs and macroscopic lesions

One group of 4 pigs was immunized intramuscularly (IM) with 10^3^ TCID_50_/mL BeninΔA179L (Group D) and boosted with 10^4^ TCID_50_/mL of the virus on both 12- and 20-days post immunization. These pigs (Group D) and a control group (Group F) of 3 non-vaccinated pigs were challenged with parental Benin 97/1 at 42 days post immunisation with 10^4^ TCID_50_/mL by the intramuscular route.

Rectal temperatures and clinical signs were recorded daily for all pigs. After the initial immunisation and the first and second boost with BeninΔA179L, the pigs showed similar average body temperatures with no increase in temperature or other clinical signs (Figures 4A, 4C). At 3 days post challenge, all pigs in group D (BeninΔA179L) developed temperatures above 40.5^0^C and exhibited other signs typical of acute ASF including lethargy and reduced eating (Figures 4A, 4C). Clinical scores varied from 6 – 12 rising to 14 – 15. One pig was euthanised on day 4 post-challenge and the remaining 3 pigs were euthanised at 5 days post-challenge at a moderate severity humane endpoint. The group F (control) pigs displayed an increase in temperature (40.6 – 41.6^0^C) and other clinical signs typical of acute ASF by three days post-challenge and all were euthanised at the moderate severity humane endpoint four days post-challenge (Figures 4B, 4D).

**Figure 4.**
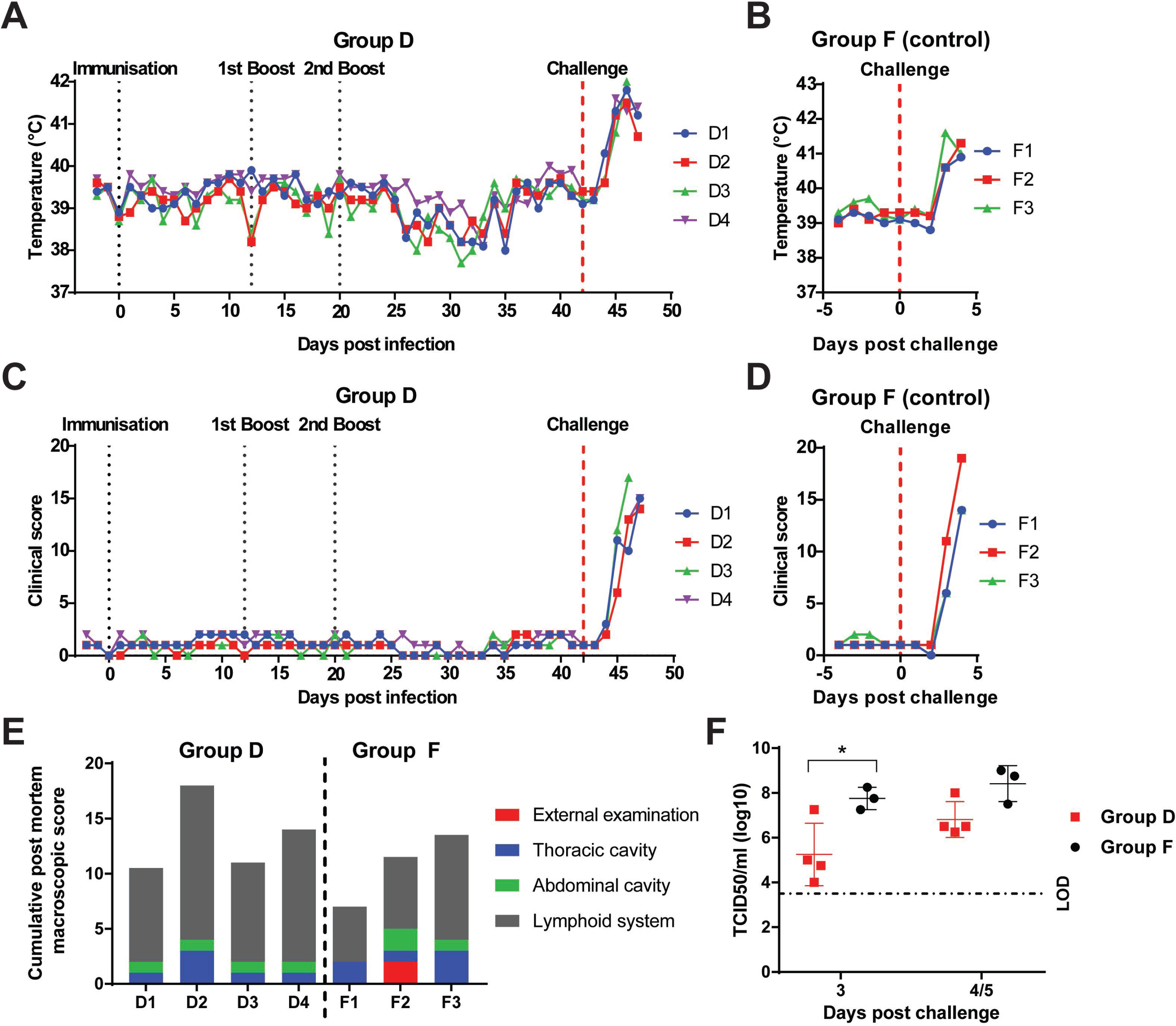
Temperatures, clinical scores, post-mortem macroscopic lesion scores and viremia in pigs immunised with BeninΔA179L and challenged with virulent Benin97/1. Rectal temperatures were recorded daily for BeninΔA179L-immunized pigs (Group D) (**A)** and non-immunized naïve control pigs (Group F) (**B**). Cumulative clinical scores based on clinical signs observed daily are shown on panel **C** (Group D) and panel **D** (Group F). (**E**) Lesions observed were scored and a cumulative score is shown for each pig. External examination (red bar) includes general body condition and conjunctiva; Lesions in the thoracic cavity (blue bar) include the presence of thoracic exudates as well as lesions affecting cardio-respiratory system. Lesions in the abdominal cavity (green bar) include the presence of ascites along with the presence of lesions affecting the gastrointestinal system including the stomach, intestines, liver and gallbladder. Finally, lesions depicted by the grey bar include pathology observed in lymphoid tissues: tonsils, thymus, spleen and various lymph nodes. (**F**) Levels of infectious virus were measured for non-vaccinated pigs (Group F) and BeninΔA179L-immunized pigs (Group D) by titration of post-challenge whole blood collected in EDTA. No virus was detected before challenge in Group D pigs. 2-way ANOVA with Sidak’s multiple comparisons test was performed to evaluate the differences between viremias in Group D and Group F. Significant differences are represented by asterisk where * is p<0.05.

All pigs in group D (BeninΔA179L) and control group F showed macroscopic lesions typical of acute ASFV at necropsy (Figure 4E). Scoring of lesions (44) showed values of between 10 and 17 in pigs in group D and 8 to 15 in group F.

### Virus levels in blood

No infectious virus was detected in blood from pigs in Group D before challenge. At 3 days post-challenge, virus was detected in blood from the pigs in Group D and the control pigs in Group F. Those in Group F displayed a significantly higher titre of virus (mean: 10^7.8^ TCID_50_/ml) relative to group D pigs (mean: 10^5.3^ TCID_50_/ml) (p= 0.0149). There was no significant difference in levels of virus in blood between the groups at the time of termination 4- or 5-days post challenge (Figure 4F).

### Immune response in immunised pigs measured by IFN-gamma ELISpot and ELISA assays

The response of PBMCs, isolated from pigs pre- and post-immunization and boost with BeninΔA179L, to stimulation with ASFV Benin 97/1 was measured by IFN-γ ELISpot assays. At the second boost, 20 days post-immunization, numbers of IFN-γ producing cells ranged between 100 – 400 IFN-γ producing cells per million cells and decreased to approximately 40 – 300 million before challenge (Figure 5A). Pigs D3 and D4 displayed a decrease in IFN-γ producing cells pre-challenge compared to the time of the second boost at 20 days post-immunization. Pig D3, which was terminated at 4 days post challenge as opposed to other pigs in group D terminated 5 days post challenge, displayed the highest levels of IFN-γ producing cells (approximately 400 IFN-γ producing cells per 10^6^ cells) at 20 days post immunization. This value decreased prior to challenge (approximately 170 spots per 10^6^ cells). In contrast, the number of IFN-γ producing cells from pigs D1 and D2 increased between the 2^nd^ boost and pre-challenge (Figure 5A).

**Figure 5.**
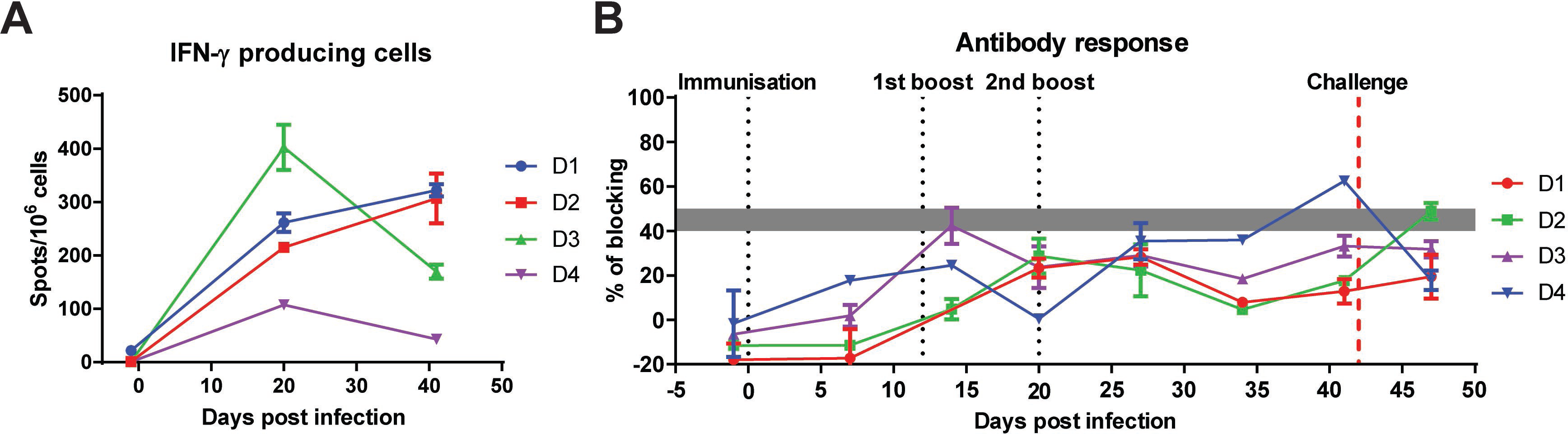
Immune responses of pigs-immunized with BeninΔA179L and challenged with virulent Benin 97/1. **(A)** The numbers of IFN-γ producing cells in PBMCs of Group D pigs were measured at 3 time points: pre-immune, pre-booster and pre-challenge by ELISpot assays. PBMCs were stimulated with Benin 97/1 isolate. Results are presented as mean frequencies of IFN-γ producing cells per million PBMC of individual pigs. (**B**) ASFV-specific antibody responses of Group D pigs were measured on different days after immunization and challenge, using a blocking ELISA against ASFV VP72. Results are presented as percentage of blocking where values above 50% blocking were considered as positive antibody responses, while anything below 40% was considered as negative. Samples with blocking between 40-50% were considered as doubtful.

A commercially available blocking ELISA assay failed to detect antibody responses to ASFV p72 capsid protein above the threshold cut-off value throughout the experiment, with exception of pig D4 just before challenge (Figure 5B).

## DISCUSSION

Host cells respond to virus infection by activating a range of defences to limit replication and induce both innate and adaptive immune responses. Signals induced by virus infection can activate apoptosis by several pathways. These include extrinsic signals, such as TNF-alpha, which binds to cell surface receptors to activate downstream apoptosis pathways. Intrinsic signals can activate pro-apoptotic BCL-2 family members leading to loss of mitochondrial outer membrane integrity, release of cytochrome *c* and initiation of caspase 3 and downstream caspase mediated cleavage. Viruses have evolved a battery of mechanisms to inhibit apoptosis. ASFV codes for several apoptosis inhibitory proteins, including a member of the IAP family of proteins A224L (22, 45), inhibitors of stress-induced apoptosis including DP71L (23, 46) protein and the BCL-2 family member A179L (21, 33, 47).

Our results confirmed that, as predicted for a BCL-2 anti-apoptosis family member, caspase 3 and 7 activity increased at early times post-infection of macrophages with the recombinant virus from which A179L gene was deleted, BeninΔA179L, compared to infections with the wild-type Benin 97/1 virus or in mock-infected macrophages. Downstream activation of effector caspases ultimately results in induction of apoptosis. We confirmed by two additional assays that the downstream activation of apoptosis was increased in cells infected with the A179L gene deletion mutant compared to wild type or mock-infected cells. In one assay we confirmed, by binding of Annexin V, that cell surface phosphatidyl serine, an early indicator of apoptosis, was increased following infection of purified PBM cells with the A179L deletion virus BeninΔA179L_mNG compared to Benin97/1_mNG. Secondly, we observed a significant increase in induction of apoptosis, as measured by DNA fragmentation using the TUNEL assay, post-infection of purified PBMs with BeninΔA179L compared to with wild type Benin 97/1. We concluded that deleting the A179L gene, reduced anti-apoptotic, pro-survival activity in infected cells and resulted in increased caspase 3 and 7 activity and downstream apoptosis. Binding of A179L to several pro-apoptotic members of the BCL-2 family has previously been described and removal of this interaction is likely to be the mechanism involved in increasing apoptosis. In previous studies (48) induction of caspase 3 was increased from early times post-infection of porcine macrophages with the attenuated NHP68 isolate compared to the virulent Lisbon 60 isolate. Similar levels of apoptosis were observed following infection with both these viruses at late times post-infection. In this study the late induction of apoptosis was also observed, although at reduced levels, in the presence of the caspase 3 inhibitor zVAD-fnk. Thus, apoptosis observed at late times post-infection was probably mediated by other pathways in addition to those induced by caspase 3 activation (49).

We also showed that deleting the A179L gene from the genome of a virulent isolate, Benin 97/1, significantly reduced virus replication in macrophages over a multi-step growth curve. However, over a single cycle growth curve, the yield of infectious virus was very similar to the wild-type Benin 97/1 virus. Surprisingly, live cell imaging, to detect infection of cells with recombinant viruses expressing a mNG fluorescent reporter, showed that the number of cells infected with the A179L gene deleted virus BeninΔA179L_mNG, compared to Benin97/1_mNG, was much higher from very early times and reached a peak by about 16 hours post-infection thereafter declining. In contrast the numbers of cells infected with Benin97/1_mNG increased more steadily reaching a plateau about 40 hours post-infection. Although a higher number of cells were observed to be infected with BeninΔA179L_mNG, similar yields of infectious virus were obtained over a period up to 16 to 20 hpi, A likely explanation is that a higher proportion of the virions in cultures infected with virus lacking the A179L gene do not reach maturity and are not infectious due to earlier onset of apoptosis. The higher number of mNeonGreen positive cells at early times post-infection with the BeninΔA179L_mNG deletion mutant virus shows a higher entry rate of virus and initiation of infection. ASFV acquires an external envelope derived from the host plasma membrane as it buds from host cells (1, 50). We observed increased Annexin V binding to cells infected with the Benin A179L_mNG deletion mutant from earlier times post-infection onward compared to those infected with Benin_mNG virus, which has an intact A179L gene. Thus, increased incorporation of phosphatidyl serine into the external ASFV envelope is predicted. Members of several enveloped virus families are known to incorporate phosphatidyl serine into their membranes resulting in enhanced cell entry by “apoptotic mimicry” mediated by binding to phosphatidyl serine receptors on the host cell membrane (51, 52, 53)). ASFV enters cells both by receptor mediated endocytosis and macropinocytosis. The latter is constitutive in macrophages and does not depend on virus specific receptors (54, 55). Possibly “apoptotic mimicry” may explain increased infection rate of cells infected with the virus lacking A179L gene. Potentially apoptotic mimicry could also extend the cellular tropism of ASFV. Further evidence is required to support these hypotheses.

The reduced virus spread, we observed in PBMs infected with viruses lacking the A179L gene at a low MOI over a multi-step growth, resulted at least in part from declining cell numbers. Since a high proportion of cells were infected, the decline in cell numbers was probably due to early induction of apoptosis in the infected cells and decreased numbers of virions reaching maturity. It’s also possible that a lower release of virus from infected cells contributed to reduced spread as documented for HIV-1. At the virus budding step, HIV-1 can be trapped on the cell surface by one family of phosphatidyl serine -binding TIM family receptors and this can be counteracted by HIV Negative Factor (Nef), by inducing the internalization of these receptors (56).

Immunisation of pigs showed that deletion of the A179L gene greatly attenuated the virulent Benin 97/1 virus. No clinical signs and no viremia were observed following immunisation or boost of pigs with BeninΔA179L. Anti-p72 ASFV antibodies were not detected except for one pig which had very low levels at one time point. Low cellular immune responses were induced, as measured by IFN-γ ELispot assays, but the pigs were not protected against challenge with the parental virulent virus Benin 97/1. Significantly lower viremia was detected in pigs immunised with BeninΔA179L 3 days post-challenge, compared to the control non-immunised pigs but on later days post-challenge, no difference in levels of viremia were observed between immunised and control pigs. We assume that *in vivo,* insufficient virus replication occurred to induce an effective protective immune response.

In contrast to our results with A179L, deletion of other apoptosis inhibitors from the ASFV genome does not reduce virus replication in cells or virulence in pigs. ASFV IAP family member, A224L, is proposed to interact directly with the processed fragment of caspase 3 (22) to inhibit apoptosis downstream of caspase 3. Despite these anti-apoptotic activities, deletion of the A224L gene from a virulent isolate did not reduce virus replication in porcine macrophage cultures or the induction and magnitude of apoptosis. Deletion of this gene also did not reduce virulence in infected pigs (57).

The ASFV DP71L protein inhibits apoptosis pathways mediated by the transcription factors CHOP/ATF4 (23, 46). The impact of deleting the DP71L gene on virus virulence in pigs varies depending on the virus isolate. Thus, deletion from the E70 isolate reduced virulence in pigs whereas deletion from the Malawi genotype VIII isolate did not reduce virulence (58). Our results show that increased caspase 3 activity and induction of apoptosis in macrophages infected with the BeninΔA179L virus correlates with reduced replication and virus attenuation in pigs. However, we cannot exclude roles of other functions of A179L protein, for example, inhibition of the autophagy regulator Beclin 1. ASFV does not encode other known BCL-2 family members and thus A179L may be the only inhibitor of the mitochondrial pathway of apoptosis. This may explain why deletion of the A179L gene had a greater impact on virus replication and virulence has compared to other apoptosis inhibitors. Our results demonstrate for the first time the important role played by the A179L BCL-2 family apoptosis inhibitor during virus replication in macrophages and in pigs.

## MATERIALS AND METHODS

### Viruses and virus titration

The Benin 97/1 genotype I (genome reference AM712239) virulent field isolate has been previously described (59). The Benin 97/1 isolate was cultured in porcine bone marrow (PBM) cells in EBSS (Sigma), supplemented with 4mM Hepes, 10% heat inactivated porcine serum (BioSera, France), 100 IU/ml penicillin and 100 μg/ml streptomycin (60). Titration of the viruses was carried out by hemadsorption assay (presented as HAD_50_/mL) or by immunofluorescence using antibodies against ASFV early protein p30/CP204L (presented as TCID_50_/mL), calculated using the Spearman and Karber formula.

### Deletion of A179L gene from ASFV isolate Benin 97/1

A transfer vector was constructed containing fragments flanking the A179L gene from the Benin 97/1 upstream and downstream. A reporter gene β-Glucuronidase under control of the ASFV p72 promoter was inserted between these flanking regions. The left flanking region was amplified with the primers 5’-GCGCAAGCTTCAGAGGGCAAAGATGGCTCAACCAC-3’ and 5’-GCGCGGATCCGATTTCCCACGGCGGTCAAGAGGAG-3’. The obtained DNA fragment of 534bp was inserted between the *Hin*dIII and *Bam*HI restriction sites within pcDNA3_VP72-GUS (position 47618 to 48151 within the Benin 97/1 genome). The right flanking region includes 178bp from A137R and was cloned using the primers 5’-GCGCGAATTCAGCGGCACCCTATATTTTTTTATTTAGG-3’ and 5’-GCGCCTCGAGCGGGGGTAAATAAAAGCTCC-3’. The right flanking region of 421bp (position 48535 to 48955 of the Benin 97/1 genome) was inserted between the *Eco*RI and *Xho*I restriction sites in pcDNA3_VP72-GUS. Pig alveolar macrophages (PAMs) were infected with Benin 97/1. Infected cells were transfected with the transfer plasmid using TransIT-LT1 (Mirus Bio, Madison, WI, USA). Recombinant viruses BeninΔA179L expressing the β-GUS gene were identified by incubation with 5-bromo-4-chloro-1H-indol-3-yl β-D-glucopyranosiduronic acid and purified by limiting dilution. The same flanking regions were used to produce BeninΔA179L_mNG, employing the mNeonGreen reporter to isolate recombinant viruses by single cell sorting as previously described (40). The control virus, Benin_mNG, was obtained using the same method, with the mNeonGreen reporter inserted in a non-coding area of Benin 97/1 between MGF300-1L and MGF300-2R.

### Complete genome sequencing of recombinant virus BeninΔA179L

Virus was grown in a 175ml flask of PBM cells. Virus DNA was purified from lysed virions in cell supernatants after treatment with DNAse I to degrade cellular DNA using a modification of the procedure used previously (61). DNA was extracted from the supernatant using MagAttract HMW DNA kit (Qiagen). DNA libraries were prepared using Illumina DNA prep kit (Illumina) and sequenced on the MiSeq instrument using a 600 cycle v3 cartridge. A total of 6085518 reads were obtained and 77.8% were assembled under the variant analysis workflow in SeqMan NGen (DNASTAR) when compared to ASFV Benin 97/1 isolate (Accession number: AM712239) with a median coverage of 3490.16. The comparative analysis confirmed the deletion of A179L gene from the 1bp to 383bp (positions 48152 to 48534 of Benin 97/1 genome) and the insertion of β-GUS (1812bp) under the control of ASFV VP72 promoter (5’-atttaataaaaacaataaattatttttataacattatat-3’).

### Virus Replication Kinetics in Cells

The Benin 97/1 and BeninΔA179L viruses were titrated in PBM cells from two different pigs. The cells were infected at a multiplicity of infection (MOI) of 0.01 (multistep growth curve) or MOI of 1 (single cycle growth curve). The cells were incubated at 37^0^C for 1 hour and then the inoculum was removed. After gently washing with PBS, fresh medium was added to the wells and the cells were further incubated at 37^0^C up to 48 hours post-infection for the single cycle growth curve or for 1 to 5 days for the multistep growth curve. Cells and supernatants were collected at different times post infection and subjected to 3 freeze-thaw cycles. Virus titres were determined as described above.

### Caspase3/7 activity assay

Purified PBMs, seeded at 5 x 10^6^ cells/mL in black polystyrene Nunc 96 well plates (Thermofisher), were cultured in RPMI supplemented with 10% heat inactivated fetal calf serum (FCS), 100 IU/ml penicillin, 100 μg/ml streptomycin and 100 ng/mL porcine CSF (Roslin Technologies) for 3 days. Prior to infection with the indicated virus at 0.5 MOI, the cells were loaded with 2 µM CellEvent™ Caspase-3/7 Green Detection Reagent (Thermo Fisher Scientific), according to the manufacturer’s instructions for kinetic assays and fluorescence in the live cells, indicative of caspase activity, recorded using a Synergy2 Multi-Detection Microplate Reader (BioTek) at the indicated time points post-infection.

### Annexin V apoptosis assay

Purified PBMs were seeded at 1.25 x 10^5^ cells per well in 96-well plates. After 2 days, the cells were treated with IncuCyte Annexin V Red reagent at 1:100 dilution in EBSS and then immediately mock-infected or infected with either Benin_mNG, and BeninΔA179L_mNG. Purified PBMs were also treated with 20µM of ABT-263 (ChemCruz, Santa Cruz Biotecnology), a mimetic of BH3 domains to inhibit members of the BCL-2 family, specifically BCL-2, BCL-xL and BCL-W. Cells were monitored for 48h using an IncuCyte S3 live-cell imaging system (Sartorius) in a 37°C incubator. Four image fields per well were captured every 2h and spectral unmixing was carried out as recommended, where 8% of the red fluorescence was removed from the green channel. Using the IncuCyte integrated software, the average Annexin V mean intensity was obtained over time and repeated measures two-way ANOVA, with Dunnett’s multiple comparison tests was used to compare the differences between the groups. To plot the mean fluorescent intensity (MFI) of Annexin V, raw cell-by-cell object data for the 22 hpi time point, where the ratio of Annexin V was highest between Benin and BeninΔA179L, was exported and analysed using FCS Express 7. The percentages of mNeonGreen and/or Annexin positive cells were calculated using the IncuCyte integrated software module, based on a user-defined cell-by-cell classification. In a similar manner to analysis by flow cytometry, wells containing controls ((i) negative for mNeonGreen and Annexin V; (ii) single colour controls – mNeonGreen positive only or Annexin V positive only; (iii) double positive for mNeongreen and Annexin V) were used to set up quadrants. Cells were then classified as (i) uninfected, non-apoptotic cells (mNG^-^AnnexinV^-^); (ii) uninfected, apoptotic cells (mNG^-^ AnnexinV^+^); (iii) infected, non-apoptotic cells (mNG^+^AnnexinV^-^); and (iv) infected, apoptotic cells (mNG^+^AnnexinV^+^). Repeated measures 2-way ANOVA with Šídák multiple comparison test was done to evaluate differences in the number of number infected, apoptotic cells over the total infected cells between the groups. This experiment was repeated in purified PBMs from 3 different pigs.

### TUNEL assay to measure apoptosis

Purified PBMs, seeded at 3 x 10^6^ cells/mL in chambered slides (Ibidi, Germany), were cultured in RPMI supplemented with 10% heat inactivated FCS, 100 IU/ml penicillin, 100 μg/ml streptomycin and 100 ng/mL porcine CSF for 3 days. The cells were infected with Benin 97/1 or BeninΔA179L at 0.25 MOI for 16 h. The cells were fixed in 4% paraformaldehyde and washed with PBS followed by the terminal deoxynucleotidyl transferase-dUTP nick end labeling (Click-iT™ Plus TUNEL Assay for In Situ Apoptosis Detection, Thermo Fisher Scientific), according to the manufacturer’s protocol. The cells were subsequently stained with mouse anti-P30 C18 (62) for 1 h at RT and corresponding goat anti-mouse IgG (H+L) cross-adsorbed secondary antibody, Alexa Fluor 488 (Thermo Fisher Scientific) at 1 ug/mL for 1 h at RT. Nuclei were labelled with Hoechst 33342, diluted in PBS at a final concentration of 5 µg/mL and incubated at RT for 15 min. Imaging was performed using a Leica TCS SP8 confocal microscope using a 40X oil immersion objective. Total number of cells and TUNEL positive cells counted using ImageJ 1.x (63).

### Immunization and challenge of pigs

Immunization and challenge of pigs with ASFV was carried out at The Pirbright Institute, Surrey UK, in high containment large animal facilities licensed at UK Specified Animal Pathogens Order level 4. Animal experiments were ethically reviewed and carried out under the Animals Scientific Procedures Act UK 1986 licensed by UK Home Office Project License PPL70/8852. Large White Landrace pigs of 15 to 20 kg weight were randomly assigned to group D which comprised 4 pigs that were immunised and boosted by the intramuscular route with gene-deleted virus BeninΔA179L then challenged by the intramuscular route with virulent parental virus Benin 97/1 in parallel with a group of 3 control pigs (Group F). The schedule and doses of virus used is described in Results. Rectal temperatures and other clinical signs were recorded daily from before the day of immunisation until termination. EDTA blood and serum samples were collected at different days post-immunization or challenge as indicated in Results to measure levels of virus and antibody and cellular immune responses. Animals were euthanized at a moderate severity endpoint. Macroscopic lesions were scored at post-mortem as described previously (44).

### ASFV specific antibody responses

A commercial blocking ELISA (Ingenasa PPA3 Compac) was used to measure the level of antibody responses against ASFV-p72 in serum, as per the manufacturer’s instructions. The following formula was used to calculate the percentage of blocking (PB): [(negative-control OD − sample OD) / (negative-control OD − positive-control OD)] × 100, where OD is optical density. Samples were considered positive if the PB was above the cut-off value of 50% blocking.

### IFN gamma ELISpot assays

Peripheral blood mononuclear cells (PBMC) were purified from heparinised blood using gradient centrifugation. The IFN-γ ELISpot assay have been previously described (64, 65). ELIspot plates (MAIPS4510, Millipore) were coated overnight at 4 °C with 4 µg/mL anti-porcine IFN-*γ* mAb P2F6 in 0.05 M carbonate-bicarbonate coating buffer and then washed with PBS. Cells were plated in duplicate at two different dilutions, typically 8 × 10^5^ and 4 × 10^5^ per well in RPMI supplemented with 10% FCS, 1 mM sodium pyruvate, 50 µM 2-mercaptoethanol, 100 IU/mL penicillin and 100 µg/mL streptomycin. Cells were then incubated overnight in a final volume of 200 µL with 10^5^ haemadsorption (HAD) units of Benin 97/1 or an equivalent volume of mock inoculum, or 2.5 µg/mL phytohaemagglutinin as a positive control. Cells were lysed by incubating for 5 min in water and then washed with PBS. Biotinylated anti-porcine IFN-*γ* mAb (P2C11), followed by streptavidin conjugated to alkaline phosphatase was used to visualise spots that were then counted using an ELIspot Reader System (AID). The number of spots per well was converted into the number of spots per million cells and the mean of duplicate wells plotted.

### Statistical analysis

Statistical analysis, as indicated, was performed using GraphPad Prism8 software. Two-way ANOVA followed by Sidak’s or Dunnett’s multiple comparison test was used to evaluate differences between groups.

## Acknowledgements

ACKNOWLEDGEMENTS

We are grateful to Animal Services at Pirbright for help with animal experiments and members of the ASFV team for helpful discussions. Support from Bioimaging, Bioinformatics and Dr Chris Neil is acknowledged.

